# Acoustofluidic Assembly of 3D Neurospheroids to Model Alzheimer’s Disease

**DOI:** 10.1101/2020.03.03.972299

**Authors:** Hongwei Cai, Zheng Ao, Liya Hu, Younghye Moon, Zhuhao Wu, Hui-Chen Lu, Jungsu Kim, Feng Guo

## Abstract

Neuroinflammation plays a central role in the progression of many neurodegenerative diseases such as Alzheimer’s disease, and challenges remain in modeling the complex pathological or physiological processes. Here, we report an acoustofluidic 3D cell culture device that can rapidly construct 3D neurospheroids and inflammatory microenvironments for modeling microglia-mediated neuroinflammation in Alzheimer’s disease. By incorporating a unique contactless and label-free acoustic assembly, this cell culture platform can assemble dissociated embryonic mouse brain cells into hundreds of uniform 3D neurospheroids with controlled cell numbers, composition (e.g. neurons, astrocytes, and microglia), and environmental components (e.g. amyloid-β aggregates) in hydrogel within minutes. Moreover, this platform can maintain and monitor the interaction among neurons, astrocytes, microglia, and amyloid-β aggregates in real-time for several days to weeks, after the integration of a high-throughput, time-lapse cell imaging approach. We demonstrated that our engineered 3D neurospheroids can represent the amyloid-β neurotoxicity, which is one of the main pathological features of Alzheimer’s disease. Using this method, we also investigated the microglia migratory behaviors and activation in the engineered 3D inflammatory microenvironment at a high throughput manner, which is not easy to achieve in 2D neuronal cultures or animal models. Along with the simple fabrication and setup, the acoustofluidic technology is compatible with conventional Petri dishes and well-plates, supports the fine-tuning of the cellular and environmental components of 3D neurospheroids, and enables the high-throughput cellular interaction investigation. We believe our technology may be widely used as in vitro brain models for modeling neurodegenerative diseases, discovering new drugs, and testing neurotoxicity.

## Introduction

Neuroinflammation involves complex biochemical and cellular responses of the central nervous system (CNS) to injury, infection or neurodegenerative diseases.^1^ To defend against potential harm, neuroinflammation is initiated in response to the triggering factors. However, as the disease progresses, the intrinsic immune response of the CNS can become either protective or detrimental to the progressing disease. In addition, acute inflammatory response initiated by traumatic brain injury or chronic dysfunction of the immune system can lead to onset or exacerbation of neurodegenerative diseases.^2, 3^ Microglia are the central players of neuroinflammation because they perform the primary immune surveillance and activities in the CNS, clearing cellular debris by phagocytosis as well as producing cytokines and chemokines to mediate secondary immune responses.^4, 5^ Microglia are essential in the development and maintenance of the CNS function. However, aberrant activation of microglia can trigger an acute inflammatory response and becomes chronic neuroinflammation.^2^

Neuroinflammation is prevailing in many brain diseases such as Alzheimer’s disease (AD), traumatic brain injury, Parkinson’s disease, and Huntington’s disease.^6, 7^Among these diseases, AD is one of the most costly neurodegeneration diseases to society, affecting an estimate of 50 million people worldwide.^8^ Tremendous efforts have been made to study the pathogenesis of AD and establish clinical trials of various treatments. However, the major cause of AD is still unclear, and there is no efficient clinical treatment despite high amounts of past and active research.^9^ Recent advances in neuroinflammation research provide new insights into the cause of synaptic loss in AD. It was found that the pathogenic neuron loss was closely related to the aberrant activation of microglia.^10, 11^ Thus, the investigation of the complex interaction between immune cells, neurons, and amyloid-β (Aβ) plaques may provide new insights into AD.

Extensive AD research projects and studies have been carried out using in vivo animal models and in vitro cell culture models. To date, the majority of experimental models are in vivo animal models based on transgenic mice that express human familial AD genes and spontaneously form Aβ plaques and neurofibrillary tangles.^12, 13^ However, there are significant differences between mice and humans, such as genetic and epigenetic backgrounds, which limit the mechanistic study and preclinical testing of AD.^14, 15^ Thus, there is an increasing need to develop in vitro models that can address these limitations and better mimic the microenvironment of the human brain. To date, 2D in vitro cultures have been extensively used in AD study,^16^ such as microglia migration^17, 18^ and phagocytosis assay^19^. Although 2D models are invaluable tools to understand the role of microglia, they failed to model the organized and dynamic establishment of cell-cell contacts. 3D arrangement of cellular interactions is particularly important for appropriate neuron-glia interactions, which is fundamental for the complex brain function.^20, 21^ These cellular contacts also play an important role in neuronal homeostasis, blood-brain barrier, and neuroinflammatory responses in the brain. In particular, neuroinflammation has emerged as a prominent cause of neurodegenerative conditions of AD.^22^ In addition, 3D cultures better recapitulate neuronal phenotypes in vivo, mechanical cues, and cellular signaling properties, which are key factors in determining cell morphology and differentiation.^23, 24^ Thus, 3D culture models offer better 3D environment with important cell-cell interaction and properties, there is an emerging trend towards the development of a 3D in vitro AD model.

So far, several 3D in vitro culture models have been developed to address the limitation of conventional 2D cultures, which hold exciting promise for AD study.^25–32^ For example, by using a microfluidic well array, the cultured neurospheroids displayed extensive neurite outgrowth and showed accumulation of Aβ and phosphorylated tau,^25^ which are the key hallmarks of AD. Another example is that the brain microenvironment was mimicked by introducing a constant flow to cultured neurospheroids in a microfluidic device. The neurospheroids cultured in constant flow showed better viability and neural network formation, while cell death and destruction of the neural network was induced after treating with Aβ.^26^ Moreover, a tri-culture system, consisting of human stem cell-derived neurons, astrocytes, and microglia, recapitulated the key features of AD neuroinflammatory processes.^27^ The migration and phagocytosis of microglia, as well as microglia-induced neurite degeneration and cell death, were observed. Despite these advances in modeling AD, there is still an unmet need to develop new technologies and models to better understand and study the AD etiology, especially regarding neuroinflammatory processes, with controlled cellular components and microenvironment representing those in vivo.

Acoustofluidics,^33–37^ combing acoustic waves with microfluidic or microfabricated devices for broad biomanipulation, may provide an alternative solution to generate 3D in vitro models for modeling AD neuroinflammation. This technique brings unique advantages that other techniques are lacking. First of all, acoustofluidic devices manipulate objects in a contact-free and label-free manner, because the acting forces on the objects are induced by the acoustic vibrations and acoustic streaming.^33, 38–41^ Moreover, acoustofluidic technology provides excellent biocompatibility,^42–44^ since it operates at a power intensity and frequency range that is similar to widely used medical ultrasound imaging such as prenatal imaging. Different from other label-free cell manipulation methods such as dielectrophoresis^45–47^, the acoustofluidics also preserves the cells or biological samples in their native culture medium or mixtures of native culture medium and Matrigel or other biomaterials.^48–50^ So far, different acoustofluidic designs and strategies have been developed for the spatial arrangement of cells or fabrication of complex cellular architectures. Bulk acoustic waves^51^ and voronoï tessellation^52^ were used to cluster or pattern cells. Acoustic streaming was employed to aggregate cells within the cell culture well-plates.^53, 54^ Moreover, we developed a series of surface acoustic wave (SAW)-based devices to assemble cell spheroids or aggregates within microfluidic chambers or capillaries.^55–58^ Acoustofluidic technology has been used to establish 3D cultures,^49, 51, 57^ engineer vasculatures,^59, 60^ print 3D microtissues,^41, 61, 62^ facilitate neurite outgrowth,^63^ and manipulate single-cell interaction and communications^64^. However, to fully explore the potential in modeling AD neuroinflammation, acoustofluidics need to meet most of the following requirements. (1) Typically, brain cells such as neurons, astrocytes, and glia are much more sensitive than stable cancer cell lines, which require very gentle handling and manipulation during the formation of 3D cultures. Moreover, it would be very convenient and user-friendly to have a 3D culture technique that is compatible with conventional cell culture dishes or well-plates in the conventional laboratory. (2) To mimic the physiological or pathological brain microenvironment, it is critical to maintaining the local gradient of oxygen, nutrients, and growth factors as well as other environmental components such as amyloid-β molecules and aggregates. (3) The neuroinflammation of AD is a chronic and complex process, which requires weeks to develop neuroinflammatory responses such as neurotoxicity and microglial activation. Thus, the platform that supports the long-term culture and real-time monitoring (e.g. time-lapse imaging) will be extremely important for modeling this complex event.

Herein, to address the above technical issues, we present a new acoustofluidic approach for modeling AD neuroinflammation. By coupling a bulk acoustic wave field, we can assemble hundreds of uniform 3D neurospheroids within minutes in a petri dish, which is compatible with all the laboratory setups and technologies. By precisely tuning the cell type and number, as well as Aβ aggregates, our technology can construct 3D neurospheroids and inflammatory microenvironments for modeling microglia-mediated neuroinflammation in AD. Moreover, the acoustically-assembled 3D neurospheroids can be cultured and monitored in the petri dish for a long-time period (e.g. several weeks). Using this approach, the toxic effects of Aβ aggregates to neurospheroids were investigated, and the dynamic cumulation and coverage of microglia to Aβ aggregates were observed using real-time imaging. The activation of microglia and the toxic effects of Aβ aggregates were further validated by using immunostaining and qRT-PCR. Based on the simplicity, reliability, and capability to be scale-up, we believe our acoustofluidic method may not only improve the understanding of neuroinflammatory diseases such as AD, and Parkinson’s disease, but also facilitate the study of autoimmune diseases such as multiple sclerosis, rheumatoid arthritis, and Crohn’s disease.

## Results and discussion

### Working principle

The acoustofluidic assembly device consisted of four 10 mm piezoelectric transducers (PZTs) arranged as orthogonal pairs around a 35 mm cell culture petri dish (**Figure 1a**). When acoustic waves were applied to cell suspensions, cells were aggregated into 3D spheroids to mimic the AD or healthy microenvironment in vivo, which enable the observation of the interaction between different cell types and inflammatory components (e.g., Aβ aggregates). When radiofrequency (RF) signal was applied to the PZTs, a 10 × 10 pressure node array was generated and cells were patterned in the same contribution (**Figure 1b**). The central area of the petri-dish, where the two sets standing acoustic waves interacted, contained typically 100 pressure nodes. Dissociated brain cells (e.g. neurons, astrocytes, and microglia) and Aβ aggregates were uniformly pushed into pressure nodes to form 100 clusters (**Figure 1c**). By controlling the components of the cell suspension in the petri-dish, the 3D neurospheroids were acoustically-assembled to mimic the healthy or AD brain microenvironment, respectively. Using this platform, we observed the neurotoxicity of Aβ aggregates and the interaction between microglia and Aβ aggregates (accumulation, coverage, and activation), at the single-cell resolution, in real-time for extended periods. These observations demonstrated that our acoustofluidic 3D neurospheroids enabled the formation of physiologically-relevant brain tissue-mimetic 3D structures.

**Figure 1.**
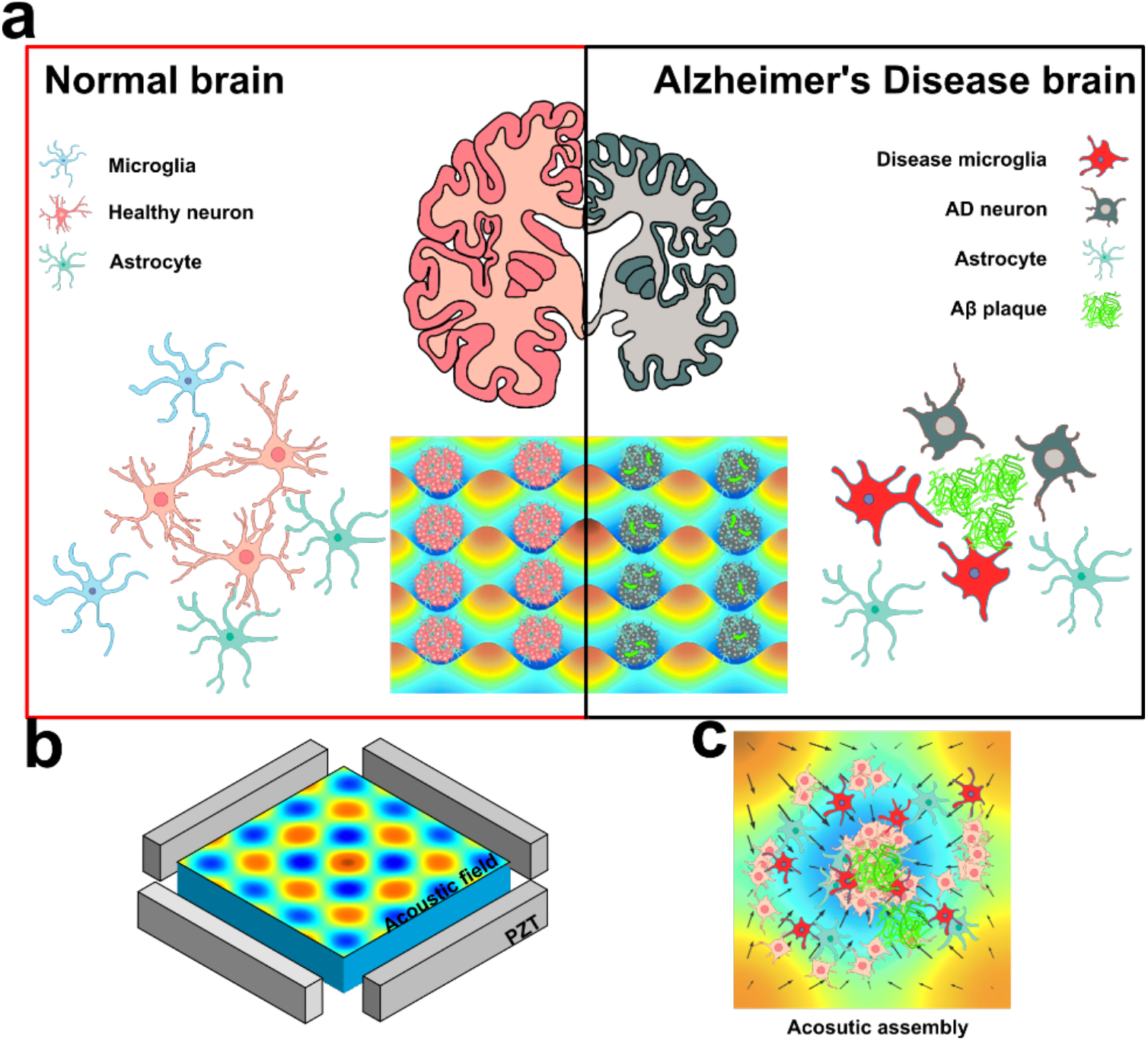
Acoustofluidic 3D cell culture to model Alzheimer’s disease. (a) Schematics of the acoustically-assembled 3D neuronal cultures to model AD. The 3D neurospheroids were generated via acoustic assembly with uniform size. By controlling the cellular and environmental components, the acoustically-assembled 3D neurospheroids can mimic the cell interaction and their microenvironment in normal and AD brain. (b) Schematic of the acoustic assembly device. The rainbow color maps the numerical simulation results of the acoustic Gor’kov potential field in the acoustic assembly chamber. Red and blue colors indicate anti-pressure and pressure nodes, respectively. (c) The acoustic assembly process of 3D neurospheroids.

### Acoustic cell assembly

We tested the capability of our acoustofluidic method for long-term culturing and maintaining uniform cell clusters. Mouse neuronal cells, Neuro 2A (N2A), were used to optimize the acoustic cell assembly of our device. N2A cells (2 × 10^6^/mL) were first introduced into the petri-dish and evenly distributed in the suspension before applying acoustic fields. Once applying RF signals at 1 MHz, two orthogonal sets of acoustic standing waves were generated. Acoustic standing waves propagated into the inner chamber, interacted with each other, and formed a periodically-distributed Gor’kov potential, which has a dot-array-like distribution, and each dot has a 3D cylinder-shaped Gor’kov potential distribution. Consequently, cells were pushed into the periodically-distributed Gor’kov potential and formed hundreds of 3D cell aggregates with the same spatial distribution (**Figure 2a, Movie S1**). These 3D cell clusters or neurospheroids were monitored every 24 hours using a fluorescence microscope. To further quantify the spatial distribution of acoustically-assembled 3D cell clusters, the images of acoustic cell patterning were analyzed and plotted along the X and Y-axis (**Figure 2b, c**). Corresponding to the brightness oscillated along the X and Y axis of defined periodicity, the 3D cell clusters were located periodically (λ/2 = 750 μm) along the X and Y axis with uniform size (163 ± 12.5 μm). The brightness curve changed sharply at the edge of the assembled clusters, indicating the capacity of generating uniform and well-defined clusters using the acoustofluidic patterning method. After a 5-day culture, the firm 3D N2A cell clusters were formed with uniformed size, while remaining in a dot-array-like pattern (**Figure 2d, e**). From the detailed view of each cluster, the 3D cell aggregates grow smooth surfaces and contained firm and complex cell-cell contact. A cell viability test was conducted to evaluate the biocompatibility of our method. The viability of N2A cells during the assembly and culture process showed no significant difference as compared to cells before acoustical assembly (**Figure 2f**). When the cell aggregates were formed and cultured in the Petri dish, high cell viability was maintained for 5 days (>90%). Thus, we demonstrated our method can generate intact and viable cell aggregates that are suitable to further model neuroinflammation.

**Figure 2.**
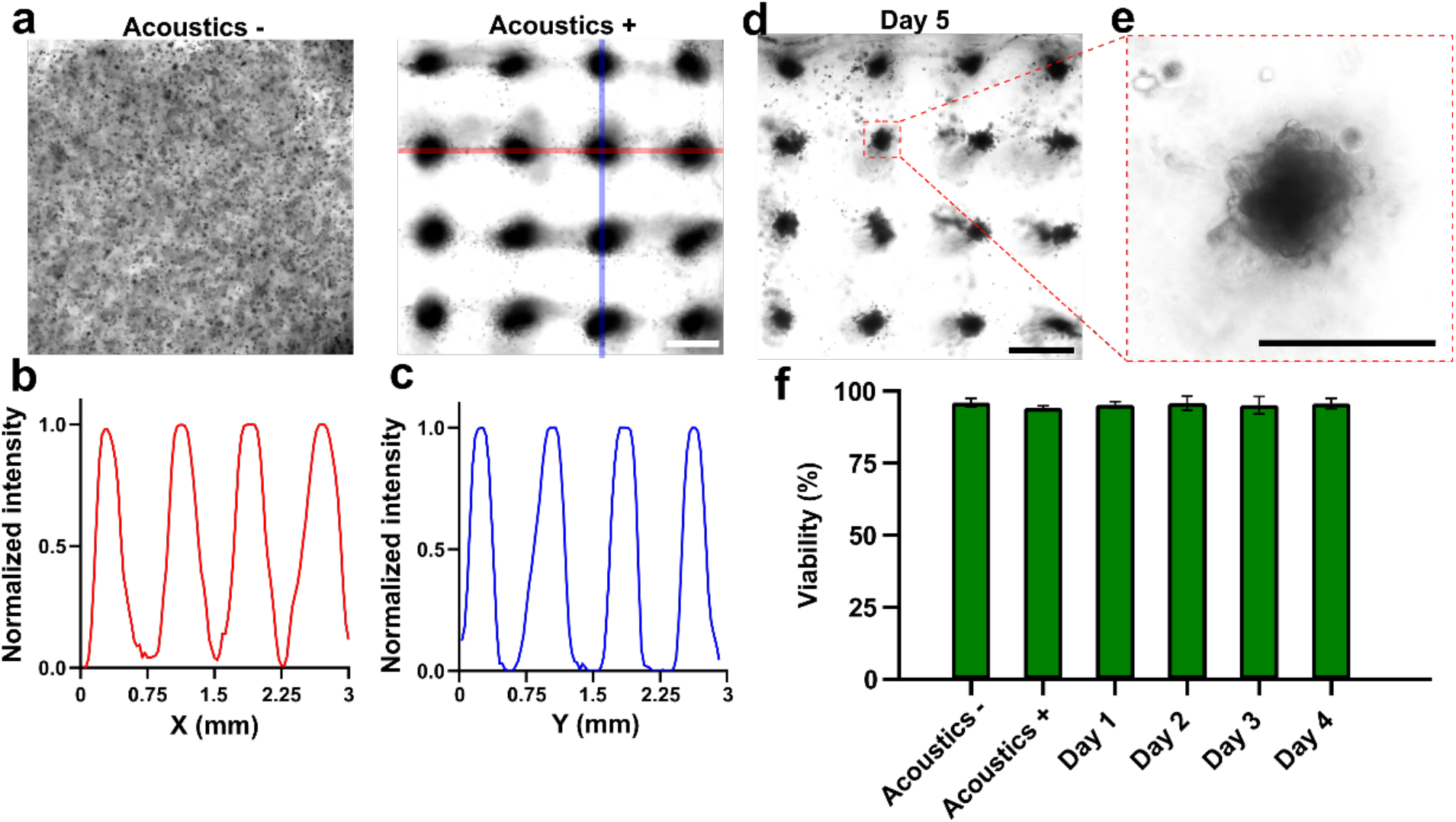
Acoustic cell clustering. (a) The acoustic assembly process of N2A cell clusters. When applied with acoustic waves, randomly distributed cells (as “Acoustics -”) migrate and form arbitrary patterned cell clusters (as “Acoustics +”, the assembling process as **Movie S-1**). (b, c) The measured brightness of pattern image along the X and Y-axis corresponding to the red and blue area in (a). The brightness result was normalized to the maximum brightness of the image. (d) Acoustic patterned N2A cell clusters after a 5-day culture. Assembled N2A cells aggregated together and formed firm neurospheroids. (e) Detailed view of single acoustic assembled N2A cell cluster after 5-day culture. (f) The cell viability was measured by LIVE/DEAD^TM^ kit, before and right after acoustic assembly, and during cell culture after the acoustic assembly. Data represent means ± s.e.m. (Scale bar = 500 μm)

### Amyloid-β toxicity

Aβ plagues or aggregates are considered as one of the key contributors to AD and they are associated with neurotoxicity and neuron dysfunction.^65^ To demonstrate the potential of acoustic methods for modeling AD, the neurotoxicity of Aβ aggregates were measured using acoustically assembled 3D neurospheroids. To explore how Aβ affects 3D neurospheroids, Aβ aggregates (5 μM) were acoustically-assembled with dissociated primary neuronal cells from an in vivo embryonic mouse brain to form cell clusters or neurospheroids with Aβ aggregates (Aβ+). The same primary neural cell suspension was also acoustically-assembled without Aβ aggregates as control groups (Aβ−). These engineered 3D neurospheroids were imaged and measured every day from day 0 (after acoustic assembly) until day 5 (**Figure 3a**). At day 0, the average size of Aβ+ and Aβ− 3D neurospheroids was similar, showing that the two groups had similar primary neuron numbers at the starting point (**Figure 3b**). During the first two days after acoustic assembly, the size of 3D neurospheroids in both groups showed an initial decrease since cells started to aggregate and form cell-cell contacts. Following initial spheroid formation, the size of Aβ− 3D neurospheroids remained unchanged in the following three days. In contrast, the spheroid size of Aβ+ 3D neurospheroids decreased over the following three days. The average size of Aβ+ neurospheroids (82.1 ± 16.3 μm) was smaller than that of Aβ− neurospheroids (121.3 ± 21.7 μm) indicating the neurotoxic effects of Aβ aggregates as the neuron death in the presence of Aβ aggregates. Thus, our 3D models demonstrated that neurotoxic effects of Aβ aggregates, which is consistent with previous reports that Aβ aggregates contribute to the neuron death in AD brain.^66^

**Figure 3.**
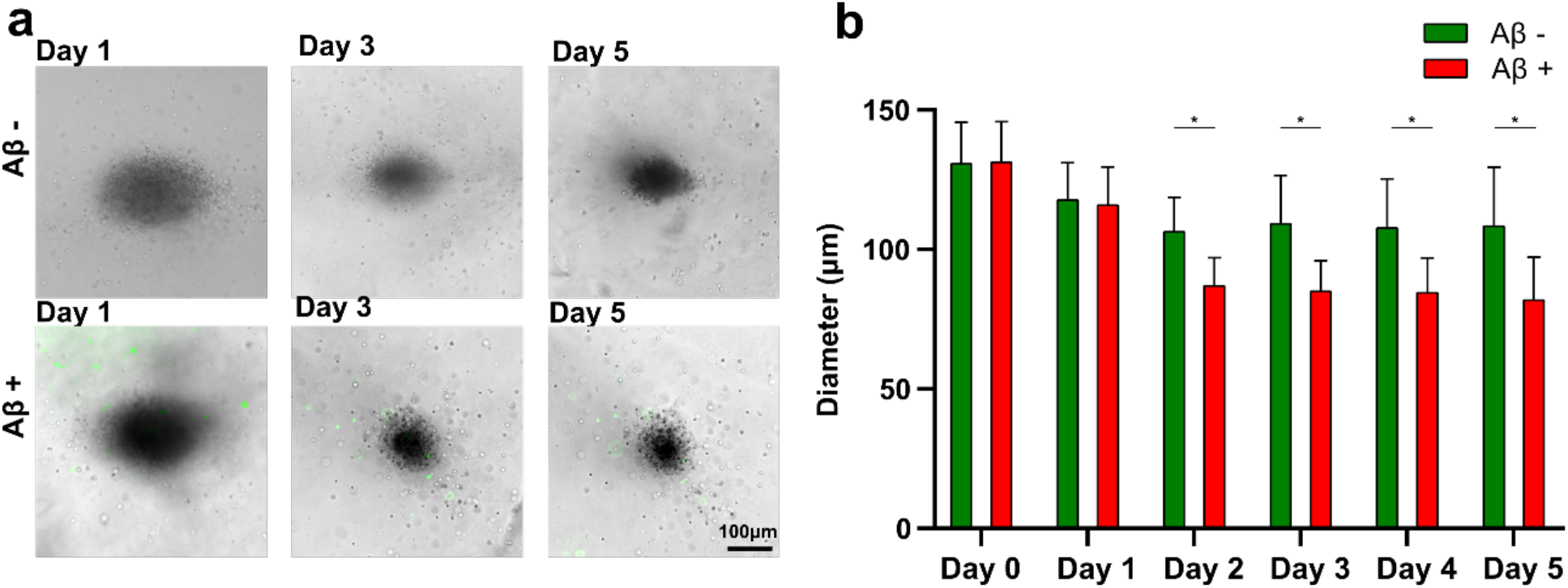
Amyloid-β Neurotoxicity. (a) Time-lapse images of primary 3D neurospheroids with or without Aβ aggregates from day 1 to day 5. The Aβ aggregates were labeled with a green fluorescent antibody against Aβ. (b) The size distribution of 3D neurospheroids with or without Aβ aggregates over time. Data represent means ± s.e.m. of 3 independent experiments (n=20, \**p* < 0.01). (Scale bar = 100)

### Model Alzheimer’s disease

Other than the neurotoxicity of Aβ aggregates, the AD brain contains more complex pathology, which is highly related to neuroinflammation.^27, 67^ The key identities associated with AD are neurons, microglia, and Aβ aggregates. To provide a more physiologically relevant system to mimic key pathological features in AD, we acoustically-assembled neurons, Aβ aggregates, and microglia together into 3D neurospheroids (**Figure 4a**). Our device can assemble randomly-distributed cellular and environmental components together into uniform 3D neurospheroids in a Petri dish, enabling a realistic model to study the complex interactions among these components. The fluorescently-labeled BV-2 microglia (Red), Aβ aggregates (Green) and primary neurons (Blue) were acoustically assembled in the trapping nodes and formed clusters (**Figure 4b**). To better mimic the in vivo conditions, we tuned the ratio of microglial cells to primary neurons inside our neurospheroids by tuning the ratio of cell suspension and finally set to be a similar ratio as in an in vivo brain (1:10).^68^ As our confocal images showed, the inner components of the 3D neurospheroids, the microglia (Red), Aβ aggregates (Green), and primary neurons (Blue) were uniformly located in the 3D neurospheroids (**Figure 4c**). These observations demonstrated that our acoustofluidic device enabled the formation of physiologically relevant 3D Aβ+ neurospheroids with the key cell types and inflammatory components.

**Figure 4.**
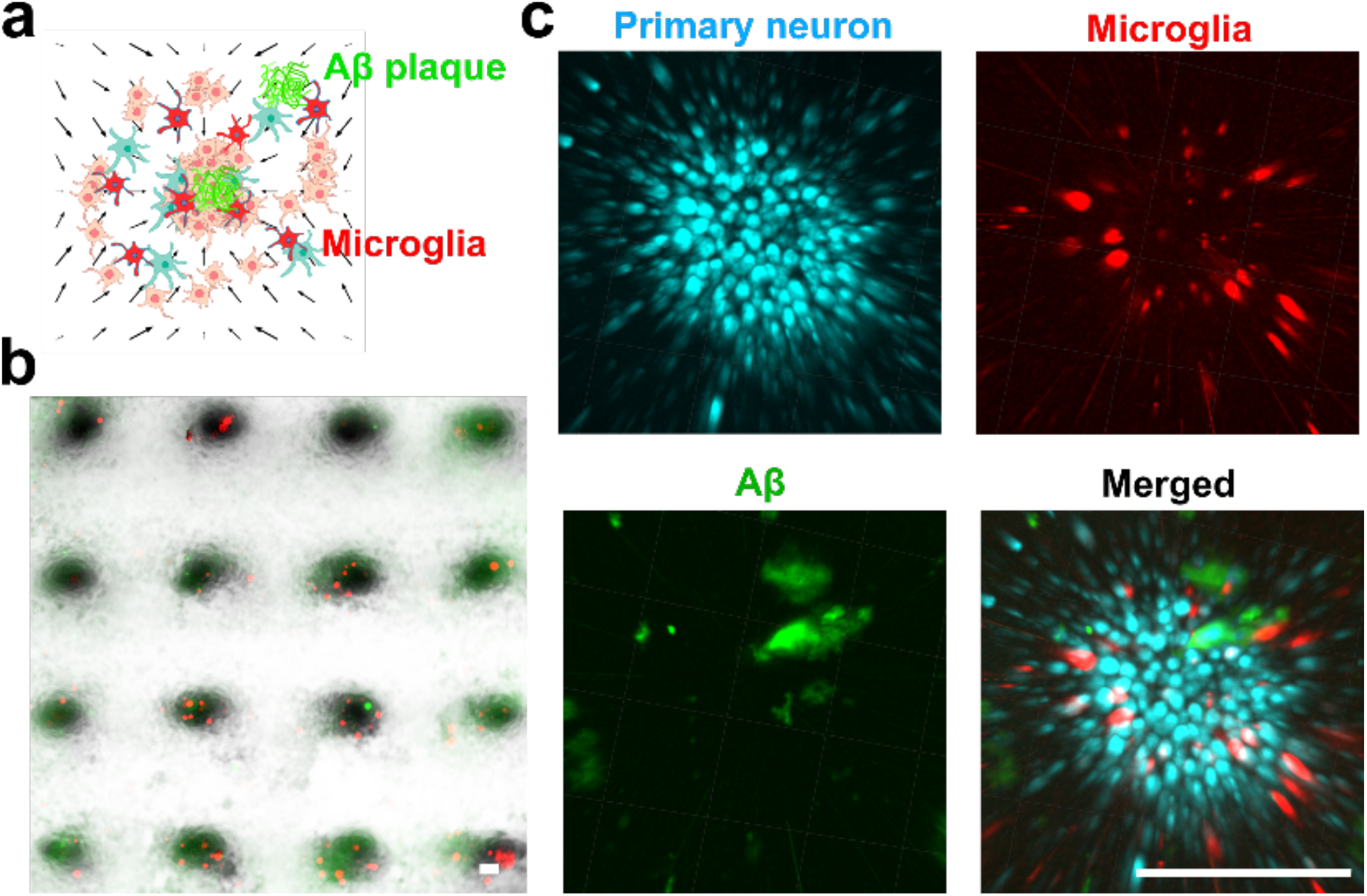
Modeling Alzheimer’s Disease. (a) Schematic of acoustically assembled 3D culture model of AD. Primary neurons (Blue), microglia (Red) and Aβ aggregates (Green) were acoustically assembled into 3D neurospheroids patterns. (b) The acoustically assembled 3D neurospheroids of AD. (c) Separate 3D reconstructed confocal images of the primary neuron (Blue) stained with CMAC dye, microglia (Red) labeled with CMTPX dye, Aβ aggregates (Green) stained with anti-Aβ 6E10 antibody, and merged images of these three colors. Microglia (Red) and Aβ aggregates (Green) were randomly distributed in the acoustically-assembled primary 3D neurospheroids (Blue). (Scale bar = 200 μm)

### Accumulation of microglia surrounding amyloid-β aggregates

In the early stage of AD, microglia migrate to Aβ plagues,^69, 70^ forming a protective barrier to protect brain tissue from the neurotoxicity of Aβ plaques, and promotes the clearance of Aβ aggregates.^71, 72^ As microglia and Aβ aggregates distributed uniformly in the acoustically assembled 3D neurospheroids, our AD model provided a realistic model for studying the accumulation of microglia around Aβ aggregates. The acoustically-assembled 3D neurospheroids were monitored using a confocal fluorescence microscope every day. The confocal images of stacks of 3D Aβ+ neurospheroids with labeled microglia (Red) and Aβ aggregates (Green) were analyzed to reveal the accumulation of microglia to Aβ aggregates. On day 0, nearly no microglia were located around the Aβ aggregates, as time went by, more microglia accumulated around the Aβ aggregates (**Figure 5a**). We further quantified the microglia accumulation to Aβ aggregates by quantifying the numbers of microglia near the Aβ aggregates (< 20 μm distance). The numbers of nearby microglia increased in the first two days up to 3 microglia per aggregate and stabilized after two days (**Figure 5b**). The microglia in the 3D Aβ+ neurospheroids accumulated to the surrounding of Aβ aggregates and the results were consistent with the previous findings in human AD brains and mouse models,^69, 72^ indicating our model provided a realistic platform to monitor the microglia accumulation in real-time.

**Figure 5.**
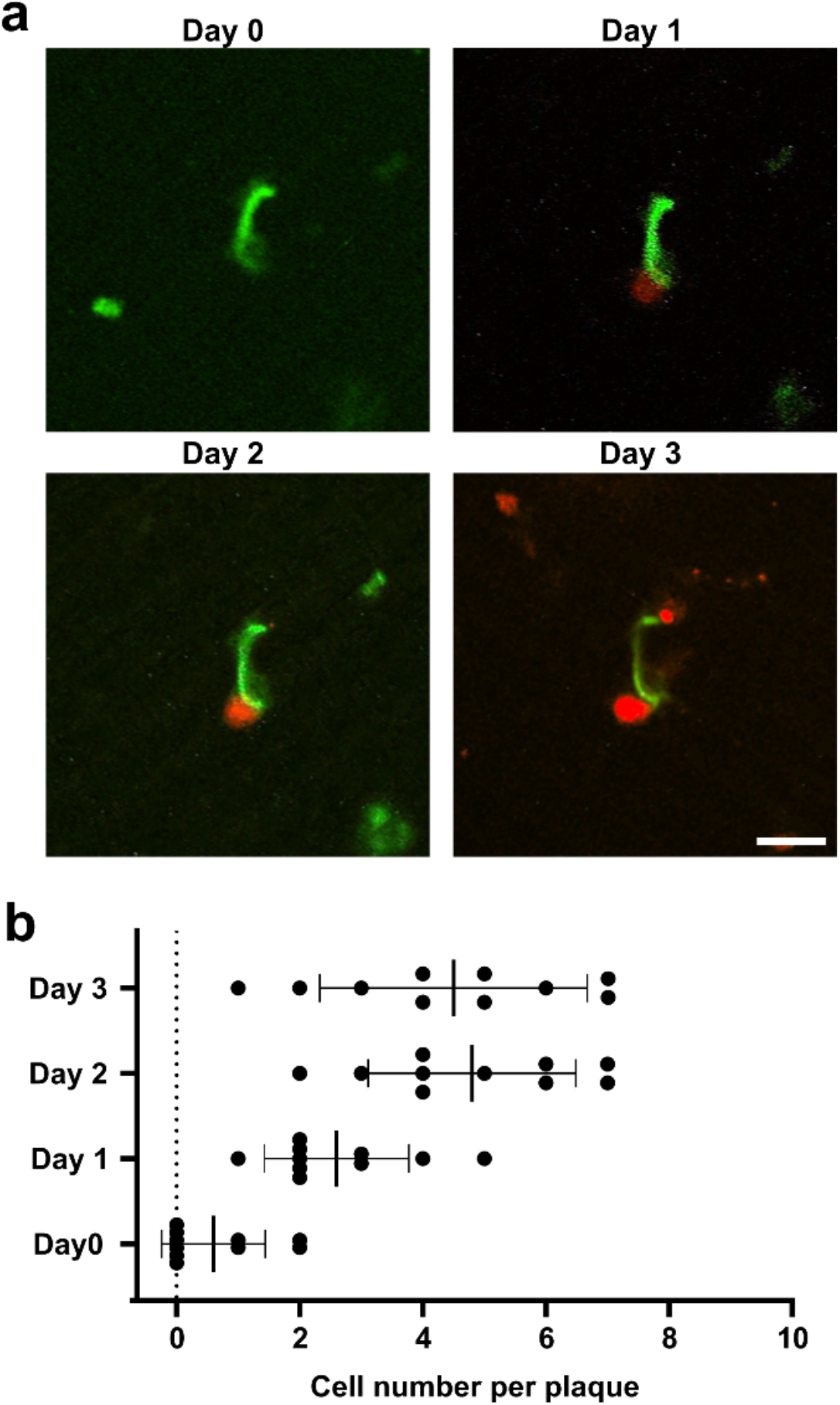
Accumulation of microglia around Amyloid-β aggregates. (a) Representative confocal time-lapse images of a red fluorescent dye (CMTPX)-labeled microglia (BV-2) around Aβ aggregates (green) stained with anti-Aβ 6E10 antibody in an acoustically assembled neurospheroid. (b) Quantification of microglia accumulation around Aβ aggregates over time. Microglia accumulation was quantified as the microglial cell numbers within 20 μm distance from Aβ aggregates. N > 10 aggregates (1 – 40 μm in diameter) from different acoustic assembled 3D neurospheroids. Bars represent mean ± s.e.m. (Scale bar = 20 μm)

### Coverage of microglia to amyloid-β aggregates

The microglia in the AD brain tightly cluster and cover around Aβ plagues and protect surrounding tissues from neurotoxicity and Aβ deposits.^73, 74^ Thus, we further analyzed the coverage of microglia to Aβ aggregates. After a 5-day culture, microglial cells accumulated to the Aβ aggregates, and clustered tightly surround those aggregates. The coverage of microglial cells to small (< 10 μm, **Figure 6a**), medium (10 ~ 20 μm, **Figure 6b**) and large (> 20 μm, **Figure 6c**) sized aggregates varied. We quantified the extent to which the surface of individual Aβ aggregates was covered by the adjacent microglia in the acoustic assembled clusters. Larger aggregates (49.3%) tended to have less microglia coverage than the smaller ones (83.2%) (**Figure 6d**). In this study, we did not observe the processes of these microglial cells, because BV-2 is a transformed microglia cell line with differed morphology compared to microglia directly isolated from the animals,^72^ which was reported previously.^75–77^ The observed relation of coverage and aggregate size was consistent with the previous in vivo study,^72^ indicating our 3D neurospheroids can recapitulate the behavior of microglia in vivo.

**Figure 6.**
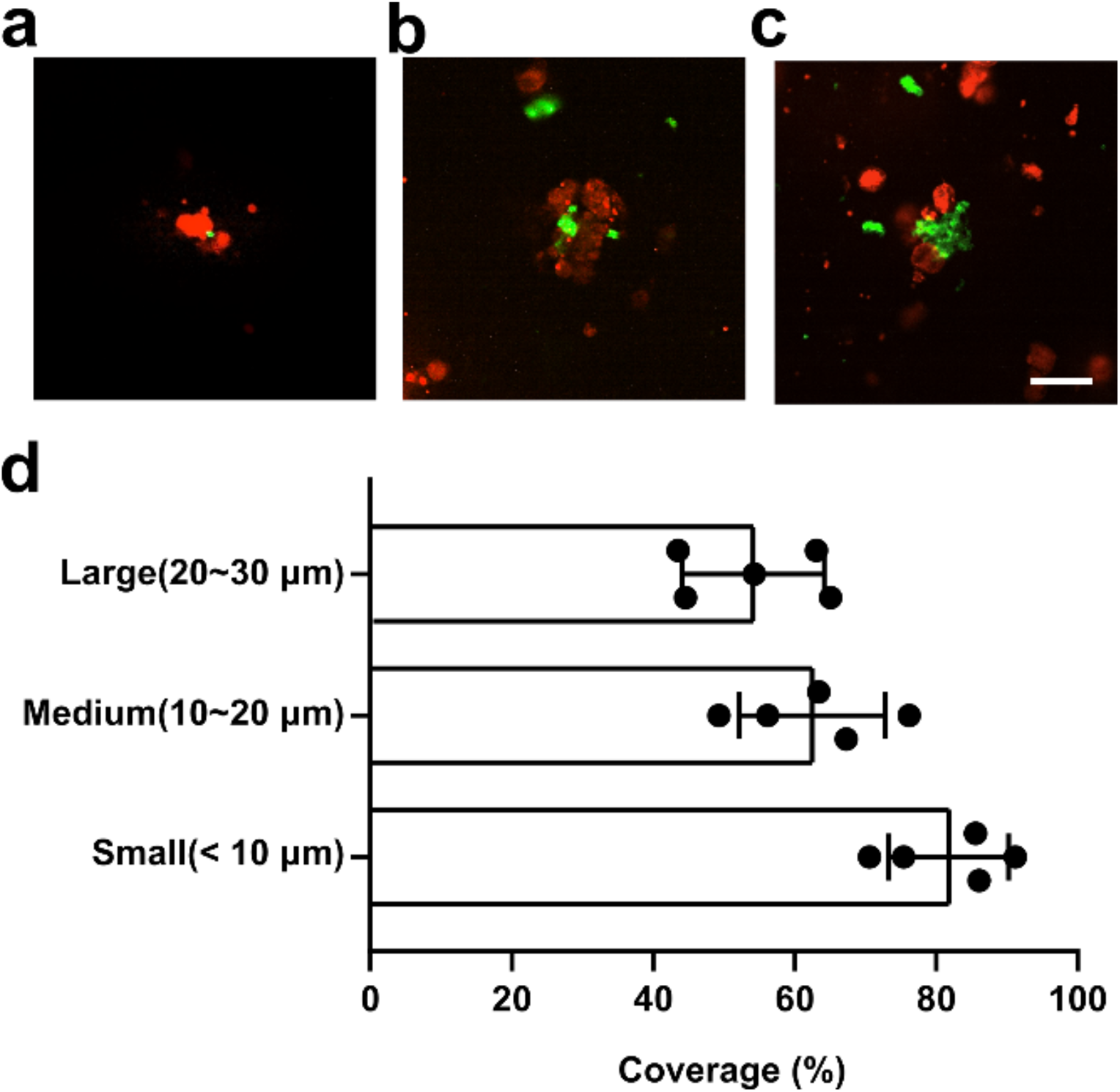
Coverage of microglia to Amyloid-β aggregates. Representative confocal time-lapse images of small (a), medium (b) and large (c) sized Aβ aggregates (green) stained with anti-Aβ 6E10 antibody covered by microglia (stained by CMTPX dye in red) in an acoustically assembled neuronal spheroid. (d) Quantification of microglia coverage in an acoustically assembled 3D neurospheroids. Microglia coverage was quantified as the percentage of aggregate perimeter contacted by the microglia process. Black bars represent mean ± s.e.m. (Scale bar = 20 μm)

### Microglia activation

In AD, brain microglial cells are activated in response to Aβ and other neuropathological changes and undergo complex neuroinflammation processes,^78^ playing either a protective or detrimental role in the disease.^79, 80^ To check the activation status of microglia in our cell culture system, 3D neurospheroids in the absence or presence of Aβ were analyzed via both immunostaining and quantitative reverse transcriptase-polymerase chain reaction (qRT-PCR). After a 5-day culture, the 3D neurospheroids with or without Aβ aggregates were immune-stained following cryo-sectioning. The 3D neurospheroids with the presence of Aβ aggregates (Thioflavin-T) expressed a higher level of ionized calcium-binding adaptor molecule 1 (Iba-1, microglia marker) and lower level of microtubule-associated protein 2 (MAP-2, neuron marker) than those without Aβ aggregates (**Figure 7a, b**), indicating the Aβ aggregates activated the microglia and may induce the neurotoxicity as shown in **Figure 3 b**. The qRT-PCR results of neuron marker NeuN and microglia marker Iba-1 also showed the same corresponding to the immunostaining results (**Figure 7c**). The Iba-1 expression in our 3D Aβ+ neurospheroids was about 7 folds higher than that in the 3D Aβ− neurospheroids. The upregulated expression of Iba-1, indicating activation of microglia, were consistent with the previous finding in vivo.^81, 82^ Taken together, our engineered 3D neurospheroids modeled the neuroinflammation such as the activation of microglia, which may provide a realistic 3D in vitro model for AD study.

**Figure 7.**
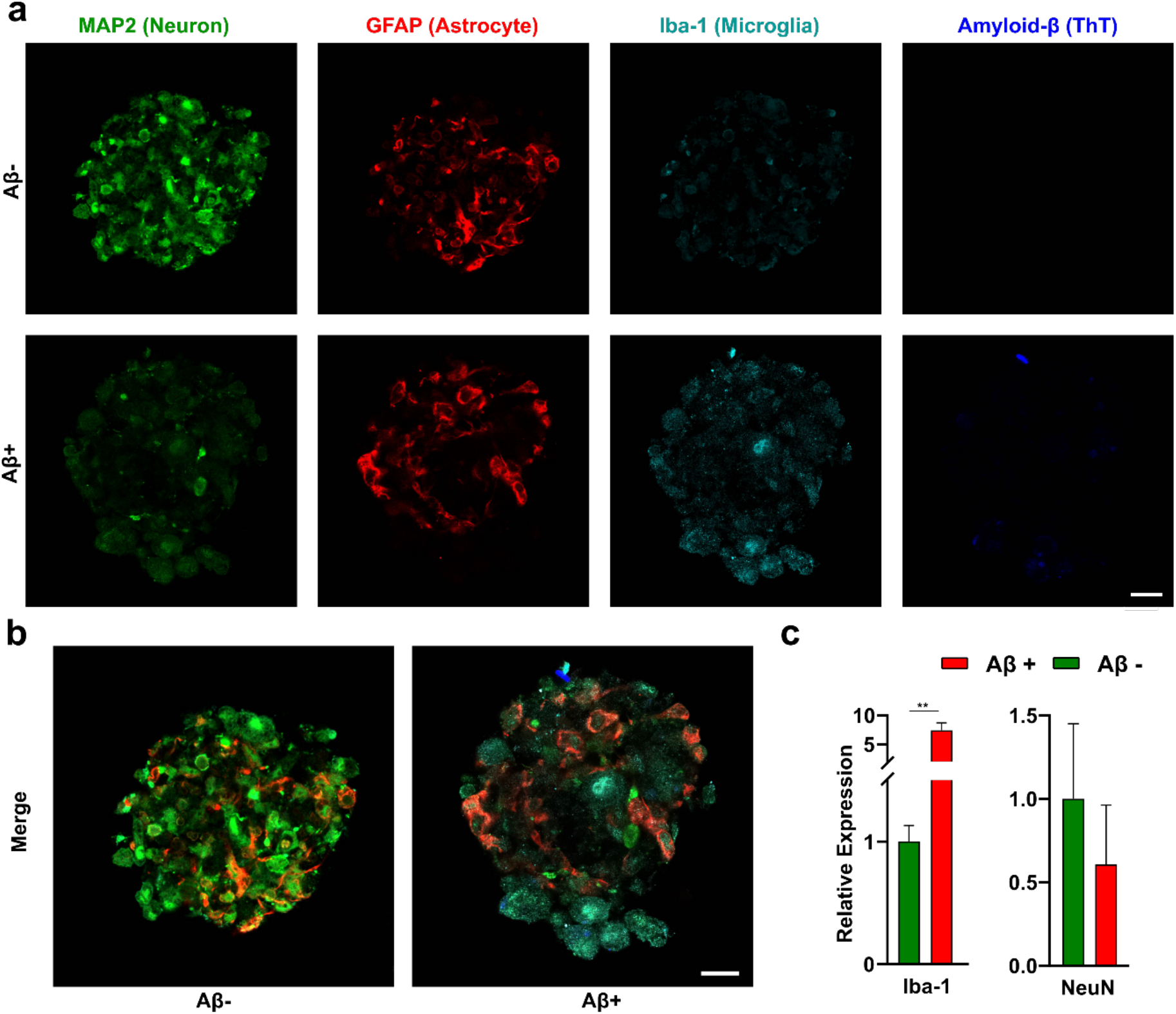
Microglia activation. (a) Representative confocal images of immune-stained acoustically assembled 3D neurospheroids after a 5-day culture without (top panel) or with (bottom panel) Aβ aggregates. (b) Merged fluorescence confocal images of neurospheroids without or with Aβ aggregates. (c) qRT-PCR results of Iba-1 and NeuN expression in acoustically assembled 3D neurospheroids after a 5-day culture without or with Aβ aggregates. Black bars represent mean ± s.e.m. (Scale bar = 20 μm)

## Conclusions

In summary, we developed a novel acoustic assembly device for high-throughput formation of uniform 3D neurospheroids using acoustofluidics. The 3D neurospheroids are assembled in the petri-dish with good viability by a device that consists of two pairs of opposite PZTs. This low-cost platform allows us to assemble 3D neurospheroids with uniformly distributed cell identities and immune components. We also used this system to investigate the neurotoxic effects of Aβ, demonstrating decreased cell viability and increased neurotoxicity, which are the key pathophysiological features of AD in vivo. Moreover, we employed this platform to study the coverage of Aβ aggregates by microglia, demonstrating the migration of microglia to Aβ aggregates, as observed in vivo brain. Our 3D culture-based microfluidic chip established the in vivo-like brain microenvironment. Therefore, it could fill the gap between traditional in vitro neuronal cell culture models and in vivo brain studies, serving as a more reliable tool for studying neurologic disease pathology and treatment strategies as well as drug screening applications.

## Experiments

### Device design and fabrication

The acoustic assembly device consists of four PZTs embedded into a laser-cut substrate and a cell culture dish. A 9 mm thick acrylic sheet was laser cut into the substrate of the device with an inner chamber of 40mm × 40mm and four small outer chambers for four embedded PZTs. The PZTs (dimension, 20 mm × 10 mm × 3mm; resonant frequency, 1MHz) were affixed to the outer chambers with epoxy, and a 3 mm thick acrylic sheet was glued to the substrate bottom to allow the chamber to contain DI water. The opposite two PZTs were wired together to a pair, and two pairs of PZTs were driven independently by two unsynchronized 1 MHz RF signals. The RF signals were generated by a function generator (TGP3152, Aim TTi) and amplified by a power amplifier (LZY-22+, Minicircuit) to drive the acoustic assembly device. A cell culture dish (35mm, Greiner Bio) was employed to contain cell suspensions and avoid contamination during the acoustic assembly process, the water-filled acrylic cavity was used to guide acoustic wave into the petri dish.

### Experiment operation

In the acoustic assembly experiment, cell suspension (2 × 10^6^ /mL) in phosphate-buffered saline (PBS) supplied with 5% Gel-MA (Sigma-Aldrich) and 1% Irgacure D-2959 (Sigma-Aldrich) were introduced into the acoustic pattern chamber. RF signals (1MHz, 10 to 25 Vpp) were applied to the PZTs to generate acoustic trapping patterns. After a 2-minute acoustic patterning, the solution was crosslinked for 30 seconds using ultraviolet light (365 nm, 6 m W/m^2^). The crosslinked solution containing 3D neurospheroids was transferred to a glass-bottom 24-well plate (MatTek Corporation) for confocal imaging and cultured in the corresponding culture medium.

### Cell culture

Neuro 2A (N2A) cells were cultured in Dulbecco’s Modified Eagle’s Medium (Corning, NY) supplemented with 10% fetal bovine serum (Sigma Aldrich, MO), 2mM GlutaMAX-1 (Gibco), 100 U/mL penicillin and 100 μg/mL streptomycin (Invitrogen, PA). BV-2 microglial cells were cultured in Dulbecco’s Modified Eagle’s Medium (Corning, NY) supplemented with 2% fetal bovine serum (Sigma Aldrich, MO), 100 U/mL penicillin and 100 μg/mL streptomycin (Invitrogen, PA). All the cells were maintained in a humidified incubator at 5 % CO_2_ and 37 °C.

### Primary neuron culture

Primary neurons were isolated from cerebral regions of untimed (around E18) embryonic CD1 fetal mice (Envigo) using a surgical procedure approved by the Institutional Animal Care and Use Committee (IACUC) of Indiana University Bloomington. Cerebral regions were dissociated into cell suspension using the Papain dissociation system (Worthington Biochemical Corporation) following the manufacture’s instruction. Primary neurons were maintained in Neurobasal medium containing B27 supplement [1 ml/ 50 ml], 0.5 Mm Glutamine Solution, 25 μM Glutamate (MW 147.13 g/Mol), and 1% antibiotic solution containing 10 000 units penicillin (Gibco) and streptomycin.

### Amyloid-β aggregates preparation

Synthetic Aβ (BioLegend) was dissolved to 1mM in 100% HFIP, aliquoted and evaporated in Nitrogen gas. The aliquots were stored at −80 C° before use. For Aβ aggregates preparation, the peptide is first resuspended in dry DMSO to 5 mM. PBS was added to bring the peptide to a final concentration of 100 μM, and shake the solution for 24 hours at 37 °C.

### Cell viability assay

The live/dead staining was conducted using a LIVE/DEAD™ kit (Invitrogen) following the manufacture’s instruction. The 3D neurospheroids were stained in medium supplemented with 2 μM of Carboxyfluorescein succinimidyl ester (CFSE) and 4 μM of ethidium homodimer (EthD) for 4 hours. And the 3D neurospheroids were washed twice and replaced with a fresh medium. The staining results were visualized by an inverted fluorescence microscope (IX81, Olympus). Final cell viability was analyzed using ImageJ to account for the area of live/dead cells.

### Label of amyloid-β and microglia

The prepared Aβ aggregates were stained with anti-Amyloid β (1:200, 6E10, Alexa 488, Biolegend) for 30 minutes prior to our acoustic assembly experiment. The BV-2 microglial cells were incubated in the serum-free culture medium supplied with red CMTPX dye (1:1000, CellTracker™, Invitrogen) for 30 minutes. The labeled BV-2 microglial cells were washed with fresh culture medium for 3 times prior to our acoustic assembly experiment.

### Immunofluorescent staining

After 5 days of culture, the 3D neurospheroids were analyzed for neuronal and neural progenitor markers using immunostaining following cryo-sectioning. Brain organoids were washed three times with phosphate-buffered saline buffer (PBS) and fixed in 4% paraformaldehyde (in PBS) at 4°C overnight. Fixed organoids were then cryoprotected in 30% sucrose overnight at 4°C. Cryoprotected organoids were embedded in cryomolds (Sakura Finetek) with O.C.T compound (Fisher Healthcare) on dry ice. Embedded neurospheroids were sectioned on a cryostat to 20μm thickness slices. Spheroid slices were then incubated with corresponding primary antibodies at 4°C overnight. Respectively, slices were stained with anti-GFAP (1:500, BioLegend), anti-Iba1 (1:200, Biolegend) and anti-MAP2 (1:500, Millipore). After primary antibody incubation, corresponding secondary antibodies (Invitrogen, 1:500) were introduced, followed by Thioflavin-T staining. The neurospheroid slices were incubated in a solution of 0.5% of thioflavin T in 0.1 N HCl for 15 minutes. The staining results were viewed using a fluorescent confocal microscope (SP8, Leica).

### Quantitative real-time PCR (qRT-PCR)

Neurospheroids were collected and lysed using RNeasy plus mini kit (Qiagen). The extracted RNA was then reverse-transcribed into complementary DNA (cDNA) using the qScript cDNA Synthesis Kit (Quantabio). Then gene expression of NeuN and IBA1 was then analyzed by SYBR green-based qRT-PCR (Life technologies). The primer sequences are as follow: NeuN forward: 5’-CCACTGAGGGAGACAAGAATA-3’, NeuNreverse:5’-AATTGCTGCAGAGACAGAGA-3’, IBA-1 forward: 5’-TGAGGAGCCATGAGCCAAAG-3’, IBA1 reverse: 5’-GCTTCAAGTTTGGACGGCAG-3’

### Statistical analysis

Data presented are quantified from at least three independent experiments. Student’s t-test was employed to determine the statistical significance (*p < 0.05, **p < 0.01, ***p < 0.001) of experiment groups. All values are presented as mean ± standard error of the mean (s.e.m).

## Supporting information

Movie-S1 Acoustic cell assembly

## Acknowledgment

This project was funded by the departmental start-up funds of Indiana University Bloomington and partially supported by the National Science Foundation (CCF-1909509), NIH awards (RF1AG056296, R01AG054102, R01AG053500, R01AG053242, and R21AG050804). The authors thank the Indiana University Imaging Center (NIH1S10OD024988-01) and Nanoscale Characterization Facility for use of their instruments.

